# Analysis of microalgae autofluorescence using full-spectrum cytometry to discriminate and monitor microalgae and bacteria

**DOI:** 10.1101/2025.07.21.666045

**Authors:** Ayagoz Meirkhanova, Sabina Marks, Damir Ussibaliev, Aizada Bexeitova, Stella A. Berger, Michael Melkonian, Ivan A. Vorobjev, Natasha S. Barteneva

## Abstract

The autofluorescence of algal pigments allows for non-invasive, high-throughput characterization of microalgae at a single-cell resolution. We applied full-spectrum cytometry, imaging flow cytometry, and cell sorting to analyze the spectral and morphological diversity among major microalgal groups and 102 Chlorophyta strains. The distinct spectral signatures from chlorophylls, carotenoids, and phycobiliproteins enabled us to achieve taxonomic resolution with clear separation of phycobiliprotein-containing taxa. Furthermore, principal component analysis of Volvocales revealed three spectral clusters supported by corresponding differences in cell size and shape. Additionally, in Gonium cultures, we observed that spectral signatures in the yellow-green region were altered by bacterial presence, suggesting that interactions between algal host and bacteria affect pigment-related fluorescence. Spectral heterogeneity observed within monocultures was linked to pigment accumulation, cell size, and morphological variability. These findings establish full-spectrum cytometry as a powerful method for profiling pigment composition, physiology, and structural diversity in microalgae, with broad applications in microbial ecology, environmental monitoring, and biotechnology.

## Introduction

Microalgae play a crucial role in aquatic ecosystems, contributing to primary production, biogeochemical cycling, and food web dynamics^1^. Pigment-based classification of microalgae is limited by the fact that different taxa often exhibit overlapping pigment profiles^2^. Chlorophylls are essential components of the light-harvesting complexes in planktic photoautotrophs, as they capture light and convert it into chemical energy. Accessory photosynthetic pigments, such as carotenoids and phycobiliproteins, expand the range of absorbed wavelengths, thereby optimizing photosynthetic performance across various environmental conditions^3^. Beyond their physiological significance, pigment data have become an established way of characterizing algal groups^2–4^. The optical properties of algal cells depend on factors like the wavelength-dependent refractive index, shape, and cell size^5,6^. However, environmental factors - such as irradiance^7–9^, spectral characteristics of light^10,11^, nutrient status^8,12^, temperature, and growth phase^13^ strongly influence microalgal pigment composition and important for industrially relevant microalgae. Still, a fundamental challenge in microalgal research remains the accurate assessment of pigment composition, taxonomic diversity, and physiological states at the single-cell level.

Traditional pigment analysis techniques, such as high-performance liquid chromatography (HPLC)^14^, delayed fluorescence spectroscopy^15^, spectrofluorimetry, and, more recently, Raman spectroscopy, have been employed for the characterization and quantification of algal pigments, offering high specificity and sensitivity ^16–19^. However, they have several limitations, including extensive sample preparation, difficulties distinguishing between dissolved organic matter (DOM) and planktonic particles, and the lack of single-cell resolution, averaging pigment content across microalgal populations^17,20^. Conventional flow cytometry (FC) is limited in terms of fluorescence detection channels^21–23^ and lacks spectral resolution^24^. While FC can differentiate broad microalgal groups, it struggles to identify genera and species. Fluorescence signals, alone or in conjunction with light scattering parameters, are often insufficient to distinguish taxonomically or functionally distinct microalgal populations due to spectral overlap^25^. Furthermore, cellular autofluorescence and optical cell parameters can vary with metabolic activity and pigment composition, complicating interpretation^5,26^. In contrast to conventional FC, which depends on predefined fluorescence channels, spectral flow cytometry (SFC) provides full emission spectral data. This enables the detection of subtle differences in spectral signatures, the resolution of closely related species within a population, and quantitative analysis of heterogenous populations^27–29^.

The objective of this study was to investigate the applicability of SFC in analyzing the pigment composition of microalgae, assessing taxonomic differentiation, and exploring physiological variability. To achieve these goals, the following aims were established: (1) to compare the spectral signatures of major microalgal groups and evaluate the role of accessory pigments in taxonomic differentiation; (2) to examine intra-order spectral variability in Volvocales isolates; (3) to assess the relationship between spectral variation and morphology; and (4) to investigate the broader applicability of spectral cytometry for high-throughput microbial characterization. By addressing these aims, this study seeks to demonstrate how full-spectrum cytometry can serve as an effective tool for investigating microalgal diversity, identifying phenotypic spectral signatures, detecting metabolic adaptations, and enhancing taxonomic classification through high-resolution, single-cell analysis.

## Methods

### Algal cultures and cytometric analysis

Algal cultures were obtained from the Central Collection of Algal Cultures at the University of Duisburg-Essen (Germany) (132 strains) and the Leibnitz Institute of Freshwater Ecology and Inland Fisheries (Berlin, Germany) (3 strains, Supplementary Tables 1-2), and maintained in Waris-H and f/2 culture medium^30,31^ at 21 °C under a 12/12 light/dark cycle in an algae growth chamber AL-30L2 (Percival, Perry, IA, USA). *Serratia marcescens* (ATCC 4003) stock was obtained from American Type Culture Collection (ATCC, Manassass, VA, USA) and grown in tryptic soy broth using MaxQ 4000 shaker incubator (Thermo Fisher Scientific, Waltham, MA, USA) at 30° C, 120 rpm or on nutrient agar petri dishes with periodical recording OD and spectral flow parameters.

For spectral analysis, the 7-laser, 186-channel ID7000 spectral cell analyzer (Sony Biotechnology, San Jose, CA, USA) was used in standardized mode. Virtual filters (VFs) were selected as a combination of fluorescent filters capturing the algal spectra variability and applied in the analysis as described earlier^29^. For imaging flow cytometry, samples were recorded with the 4-laser ImageStreamX MarkII instrument (Amnis-Cytek, Fremont, CA, USA). The cell sorting was performed using the MA900 cell sorter (Sony Biotechnology, San Jose, CA, USA).

### Data analysis

Fluorescence intensity data extracted from constructed overlay plots as “.csv” files using ID7000 software (2.0.2.17121). Further analysis of emission intensity profiles was conducted in RStudio (vs. 2023.06.1) utilizing packages such as ggplot2 (3.5.1), tidyverse (2.0.0), factoextra (1.0.7), and vegan (2.6-4), among others. The spectral region definitions were based on known pigment emission profiles and included green-yellow (∼494-566 nm), yellow-red (∼566-647 nm), and red (∼647-712 nm) regions. Coefficients of variation (CV) were calculated across detectors and compared using the Kruskal-Wallis and Wilcoxon ranksum tests. Principal component analysis (PCA) was employed to reduce dimensionality after normalizing the data using z-scores. The resulting principal component scores were used for k-means clustering, and the assigned clusters were projected onto PCA plots to visualize the separation. To visualize spectral intensity profiles, a biexponential transformation was applied. All plots were generated using ggplot2 (vs. 3.5.1). Additionally, IFC data, including cell sizes and shapes, were analyzed using IDEAS software vs. 6.2 (Amnis-Cytek, Fremont, CA, USA).

## Results

### Phycobiliprotein-driven variability in microalgal autofluorescence

We analyzed the autofluorescence spectra of 32 representatives from nine major microalgal groups, each displaying distinct autofluorescence patterns, particularly in relation to the presence or absence of phycobiliproteins (PBPs) (Supplementary Table 1). PBPs, such as phycocyanin and phycoerythrin, are known to emit fluorescence in the 570-650nm range, while chlorophyll fluorescence is typically observed between 650-720nm^32,33^. To maintain figure clarity, a subset of representative species’ spectral overlays was included in Figure 1 a and 1; complete overlay plots can be found in Supplementary Figure 2. Our analysis recvealed a primary distinction between PBP-containing taxa (Cyanobacteria, Cryptophyta, Rhodophyta, and Glaucophyta) and non-PBP taxa (Chlorophyta, Bacillariophyta, Haptophyta, Euglenophyta, and Dinophyta). Notably, there were significant differences in the spectral profiles within the 570–600 nm range (Figure 1a and 1b), corresponding to the emissions of phycocyanin and phycoerythrin, whereas non-PBP taxa primarily exhibited fluorescence associated with chlorophyll (650–720 nm). To quantify these spectral differences, we identified the following regions of interest: R1 (∼494-566 nm), R2 (∼566-647 nm), R3 (∼647-712 nm) (Figure 1a,b), and calculated fluorescence ratios for these regions (R1/R3, R2/R3, and R2/1), as shown in Figure 1c-h.

**Figure 1.**
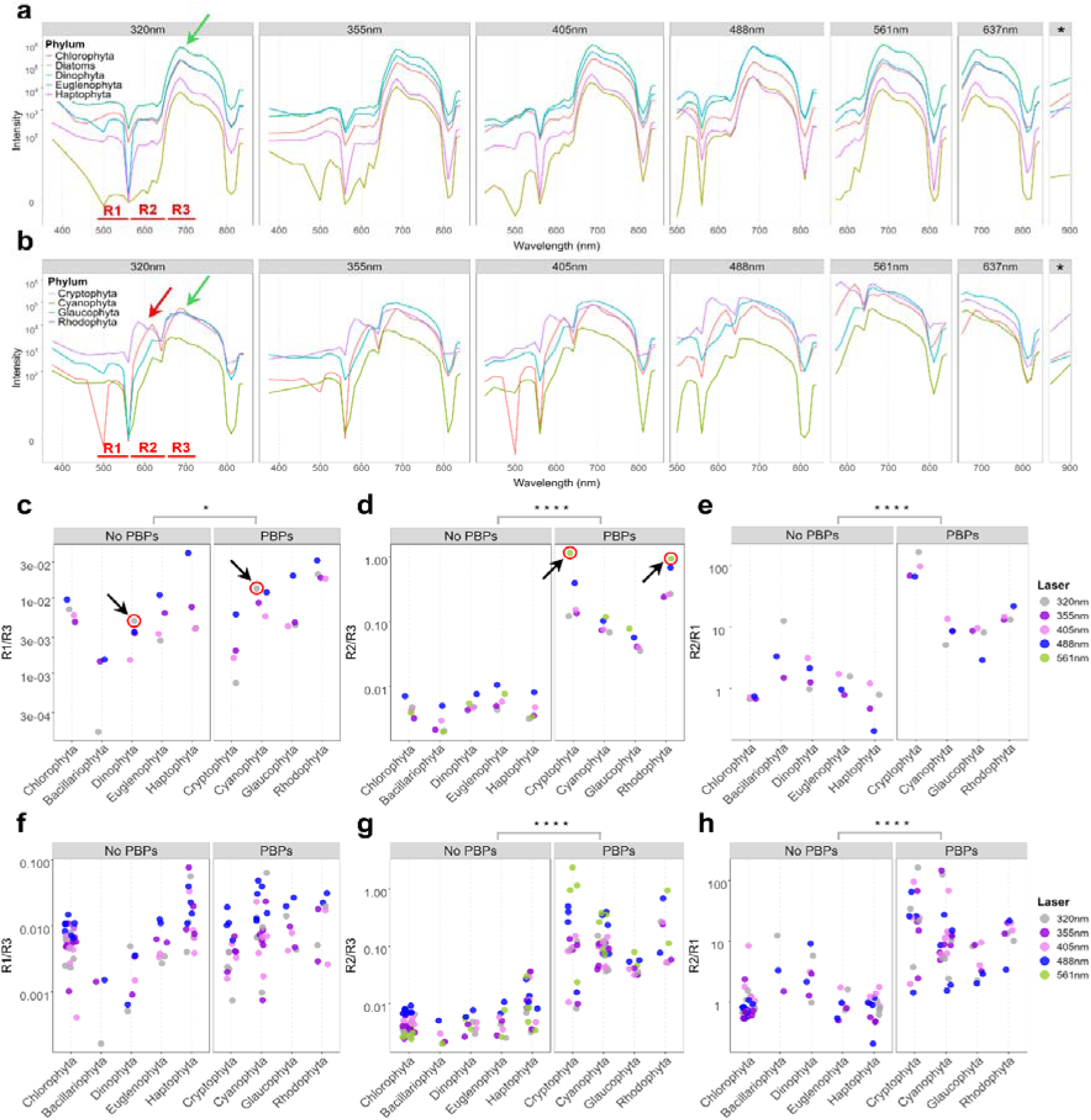
Spectral signatures and peak ratio comparisons of nine major microalgal groups with and without phycobiliproteins (PBPs); * - 808 nm excitation laser wavelength; colored arrows indicate characteristic emission peaks: green arrows mark chlorophyll-associated peaks, and red arrow indicates PBPs-associated peak (example provided for 320 nm excitation). **(a)** Overlay of spectral signatures for five representative species from microalgal groups that lack PBPs (Chlorophyta – *Chlamydomonas* sp. CCAC 3322, Bacillariophyta – *Cyclotella* sp. CCAC 3539B, Dinophyta – *Gymnodinium* sp. CCAC 3621, Euglenophyta – *Euglena sanguinea* CCAC 3518B, Haptophyta – *Prymnesium* sp.), recorded across multiple excitation wavelengths. **(b)** Overlay of spectral signatures for four representative species from PBP-containing phytoplankton groups (Cryptophyta – *Cryptomonas* sp., Cyanophyta – *Microcystis* sp., Glaucophyta - *Glaucocystis nostochinearum* CCAC 2234B, Rhodophyta – *Porphyridium cruentum* cf. CCAC 3771); Comparison of peak intensity ratios between PBP-containing and PBP-lacking groups across five excitation lasers, arrows indicate ratio values of interest: **(c)** R1/R3 ratio, **(d)** R2/R3 ratio, **(e)** R2/R1 ratio for the analysis of 9 species and 32 species **(f-h)** respectively (significant differences between groups were assessed using the Wilcoxon rank-sum test, * *p*-value<0.05, **** *p*-value<0.0001).

The R1 region exhibited significant heterogeneity in fluorescence intensity across different excitation wavelengths (320 nm, 355 nm, 405 nm, and 488 nm) (Figure 1c). Dinophyta and Cyanobacteria were distinct from other groups, exhibiting an R1/R3 maximum under 320 nm excitation. In contrast, Cryptophyta and Rhodophyta showed an R2/R3 maximum under 561 nm excitation (Figure 1c,d). The Wilcoxon rank-sum test was applied to compare the differences in the calculated ratios between PBP-containing and non-PBP groups. Our analysis of nine selected representative species revealed significant differences in the R2/R3 ratio between the two groups (p-value < 0.0001), with PBP-containing taxa exhibiting higher values (Figure 1d). Similarly, the R2/R1 and R1/R3 ratios also differed significantly between the groups (p-value < 0.0001 and p-value < 0.05, respectively). It is important to note that while the R1/R3 ratio was showed a significant difference between the two groups in the analysis of the nine species, this was not observed when examining 32 taxa (Figure 1f). The significant differences in R2/R3 and R2/1 ratios remained evident even with increasing taxonomic representation, indicating their potential as reliable discriminatory features. Additionally, the higher fluorescence intensity values in the R2 (∼566-647 nm) observed for PBP-containing taxa indicate that this spectral region may be a distinguishing characteristic for this group. To further investigate the extent of spectral variability within lower taxonomic ranks, we focused on the order Volvocales, which includes species exhibiting a range of ecological and physiological traits.

### Spectral variability in Volvocales

Based on the spectral patterns observed in major microalgal groups, we investigated the heterogeneity of spectral autofluorescence profiles within the single order Volvocales (Figure 2a). The Chlorophyta were of particular interest as a group with the reported largest variability of pigments^34^. To quantify this variability, we calculated the coefficient of variation (CV) for each detection channel, which is defined as the standard deviation divided by the mean fluorescence intensity across all 102 isolates (Figure 2b). The CVs were calculated separately for each laser, and the resulting values are presented as overlaid profiles for all lasers. As in the previous analysis, we focused on three spectral regions: R1 (approximately 494-566 nm or channels 4-9), R2 (approximately 566-647 nm or channels 10-16), and R3 (approximately 647-712 nm or channels 17-22).To determine which spectral regions exhibited greater overall variation, we grouped channels into the regions R1-R3. The comparison of CVs revealed significantly higher variability in the R1 and R2 regions compared to R3 (Kruskal-Wallis p-value < 0.001) (Figure 2c). Additionally, the emissions in the violet-orange range (channels 1–13, approximately 400–600 nm), associated with accessory pigments, displayed significantly higher variation with when excited at 561 nm, resulting in the highest degree of variation among all lasers (Figure 2b).

**Figure 2.**
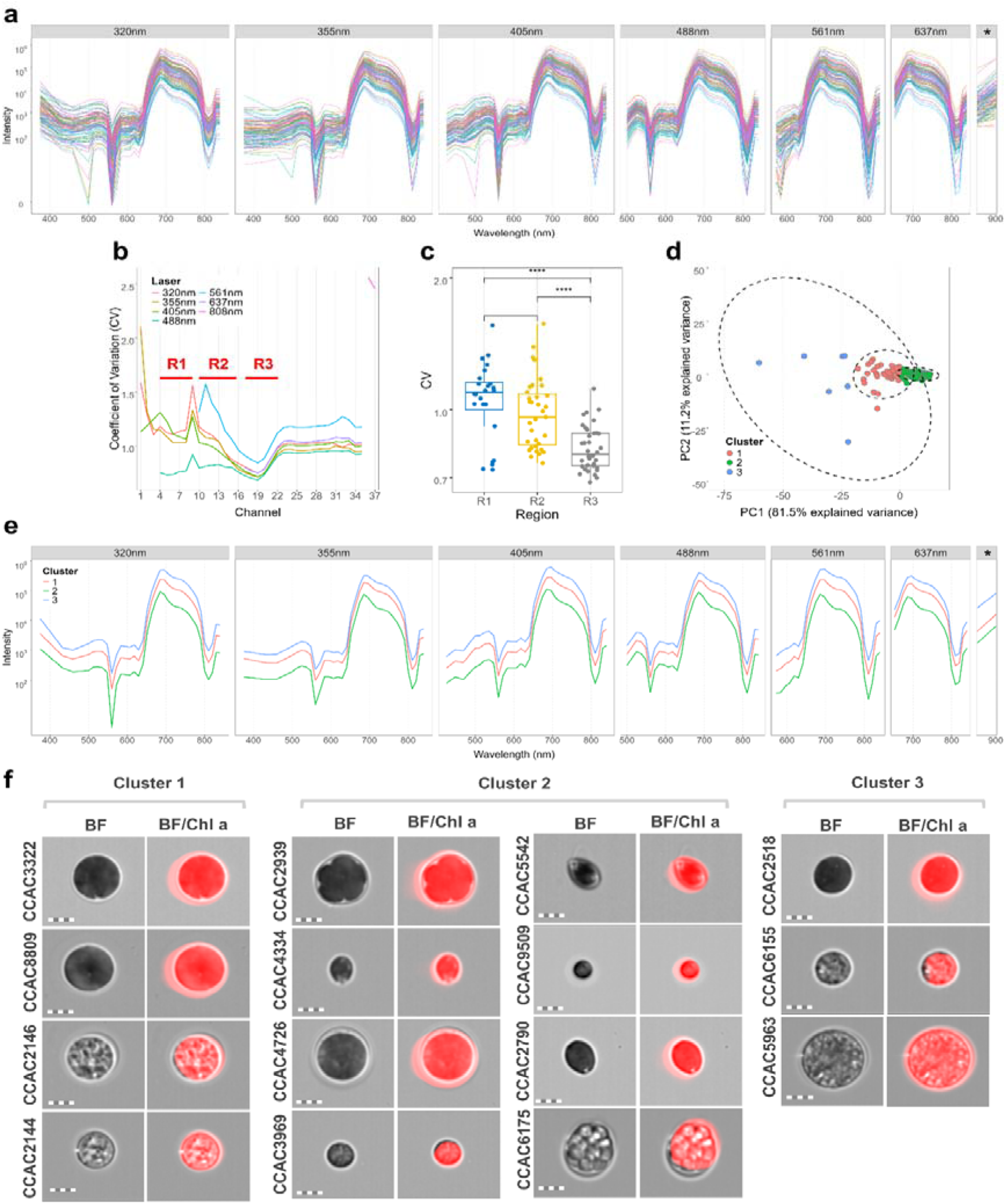
Spectral signatures and signal variation of 106 Volvocales strains, * - 808 nm excitation laser wavelength. **(a)** Overlay of autofluorescence spectra from the strains across seven excitation lasers. **(b)** Plots represent **c**oefficients of variation (CV) of fluorescence intensity across all detection channels; spectral regions (R1, R2, R3) are indicated to highlight their respective channels. **(c)** Comparison of CV distributions among regions R1, R2 and R3 (each dot represents the CV of an individual channel within the respective region), significant differences between groups were assessed using the Kruskal-Wallis test. **(d)** PCA-based kmeans clustering, revealing three distinct spectral clusters (1 - red, 2 – green, and 3 - blue) among strains. **(e)** Mean fluorescence spectra of the identified clusters. **(f)** Image library representing randomly selected strains from each of the identified clusters in brightfield (BF) and brightfield/chlorophyll *a* composite (BF/Chl *a*); scale bar = 10µm.

Principal component analysis (PCA) combined with k-means clustering revealed three distinct autofluorescent clusters, indicating intra-order heterogeneity in pigment composition and autofluorescence intensity (Figure 2d). We plotted the mean spectral signatures on the overlay to highlight the differences in spectral profiles (Figure 2e). Cluster 3 displayed the highest overall fluorescence intensity, followed by Cluster 1 and Cluster 2, particularly in the chlorophyll peak region (Clusters 1 and 3). To determine whether these differences reflect morphological variations, we analyzed 15 selected strains using imaging flow cytometry (IFC). The increased variability in the chlorophyll region could be linked to differences in cell morphology, specifically cell size and shape, as shown by IFC results (Figure 2f, Supplementary Table 3). Images of microalgae from Cluster 3 exhibited the highest fluorescence intensity and the largest average cell area (465.63 µm^2^), followed by Cluster 1 (396.38 µm^2^) and Cluster 2 (239.1 µm^2^).

### Identification of multiple autofluorescent populations within monocultures

While spectral analysis successfully identified major taxonomic groups of microalgae (Figure 1) and unique clusters within Volvocales, further investigation revealed that even single monocultures exhibited significant autofluorescence-based heterogeneity. The analysis integrated three methods: first, spectral recordings were conducted using detection channels configured to mirror the optical layout of the cell sorter MA900 (Sony Biotechnologies, San Jose, CA, USA) using the virtual filter system. This was followed by sorting of populations of interest and imaging analysis of the sorted populations using the ImageStream X Mark II (Amnis-Cytek, Fremont, CA, The results demonstrated, that in addition to genus-level spectral differences, individual Volvocales strains also contained distinct autofluorescent subpopulations linked to various cell growth phases and apoptotic cells, with characterizing by decreasing Chl *a* autofluorescence^35^. For instance, the *Chlamydomonas* sp. strain (CCAC 2145) isolate revealed up to five distinct autofluorescent subpopulations (A1, 2, B, C, and D; Figure 3b). Populations A1 and A2 exhibited similar spectral profiles with strong autofluorescence in the chlorophyll region (∼647–712 nm., However, A2 cell population showed elevated fluorescence in the 400–650 nm range. ImageStream-based IFC analysis revealed that A1 primarily consisted of single cells, while A2 was made up of dividing cells. In contrast, populations B, C, and D displayed progressively reduced chlorophyll fluorescence alongside increased emissions in the 400–550 nm and 570-640 nm regions, suggesting potential physiological stress. Notably, subpopulations differing in morphology (e.g., shape) or metabolic state (e.g., division, stress) exhibited elevated fluorescence in these regions, suggesting a potential link between fluorescence intensity and cellular physiology.

**Figure 3.**
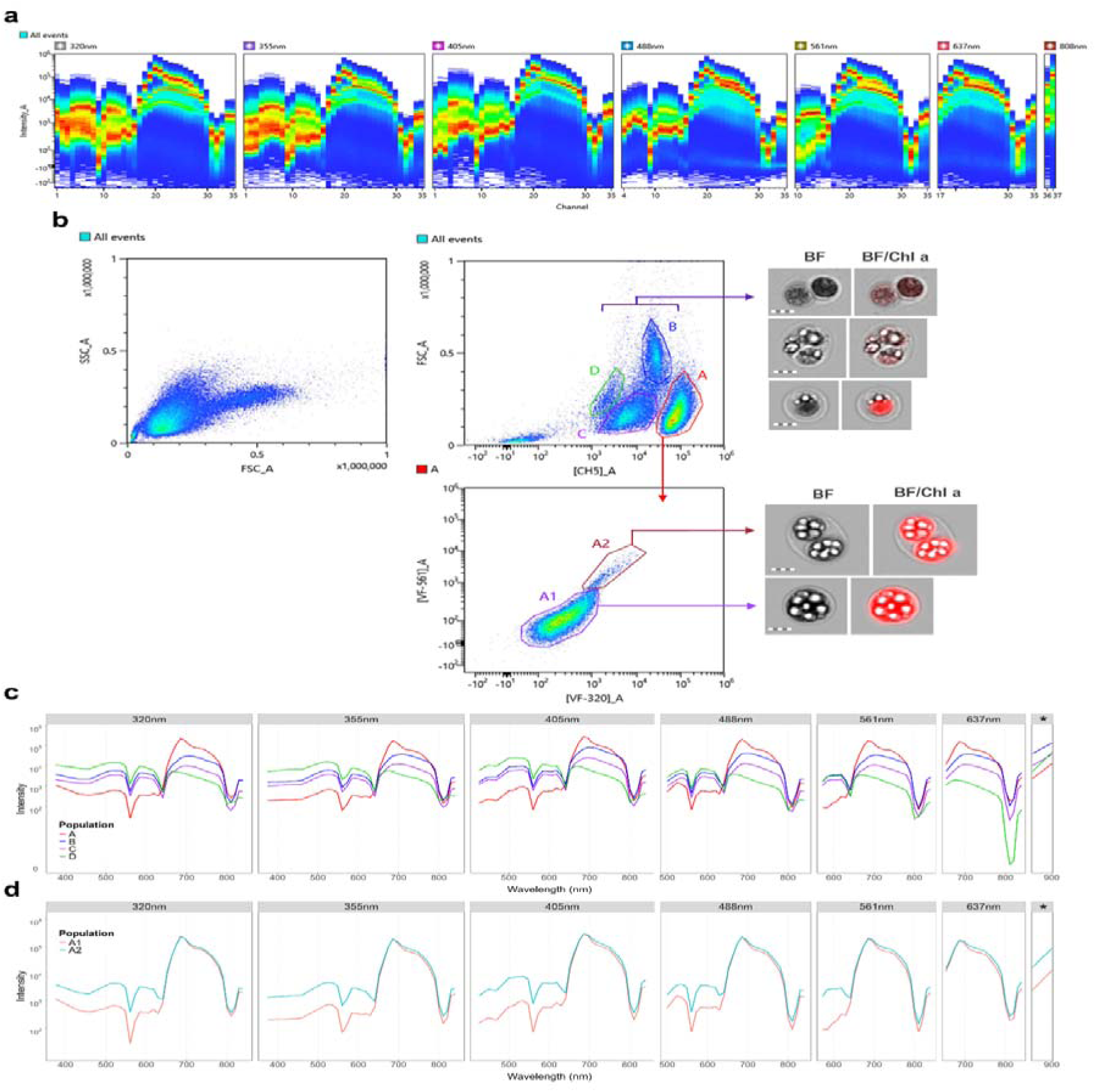
Identification of multiple autofluorescent subpopulations in *Chlamydomonas* sp. cultures, * 808 nm excitation laser wavelength. **(a)** Ribbon plot demonstrating emission signal captured from all events within the sample. **(b)** Multiple autofluorescent populations identified within individual *Chlamydomonas* sp. culture using a combination of pre-defined virtual filters (FSC_A – forward scatter area, SSC_A – side scatter area, [CH5]_A – spectral range 647-744nm under 488nm excitation, [VF-320]_A – spectral range 518-601nm under 320nm excitation, [VF-561]_A – spectral range 566-613nm under 561nm excitation) with parallel characterization of populations using imaging cytometry (scale bar = 10µm). **(c)** An overlay plot of spectral signatures of identified subpopulations. **(d)** An overlay plot of spectral signatures of A1 and A2 subpopulations.

Next, to characterize the spectral heterogeneity within *Chlamydomonas* sp. cultures, we analyzed two isolates in detail: *Chlamydomonas* sp. CCAC 9509 and CCAC 2143 (Supplementary Figures 3 and 4). For CCAC 9509, our analysis revealed two distinct autofluorescent subpopulations. The first subpopulation primarily consisted of smaller, single cells with a mean cell area of 94.01 μm² (Supplementary Table 4), while the second subpopulation comprised larger, dividing cells or doublets, averaging 208.51 μm². These two populations differed in overall emission intensity approximately 1-log) - and in spectral shape (Supplementary Figure 3). In the case of CCAC 2143, three subpopulations were detected and distinguished by cell shape: population “A” consisted of rounder cells with a circularity of 12.99 (Supplementary Table 5), while populations “B” and “C” were more elongated, with a circularity of 5.28. Spectral analysis confirmed the presence of three distinct autofluorescence profiles; however, no major differences were observed under 561 nm excitation (Supplementary Figure 4). These findings indicate that autofluorescence can reveal subtle physiological differences related to cell morphology, division state, or metabolic activity in microalgal populations.

**Figure 4.**
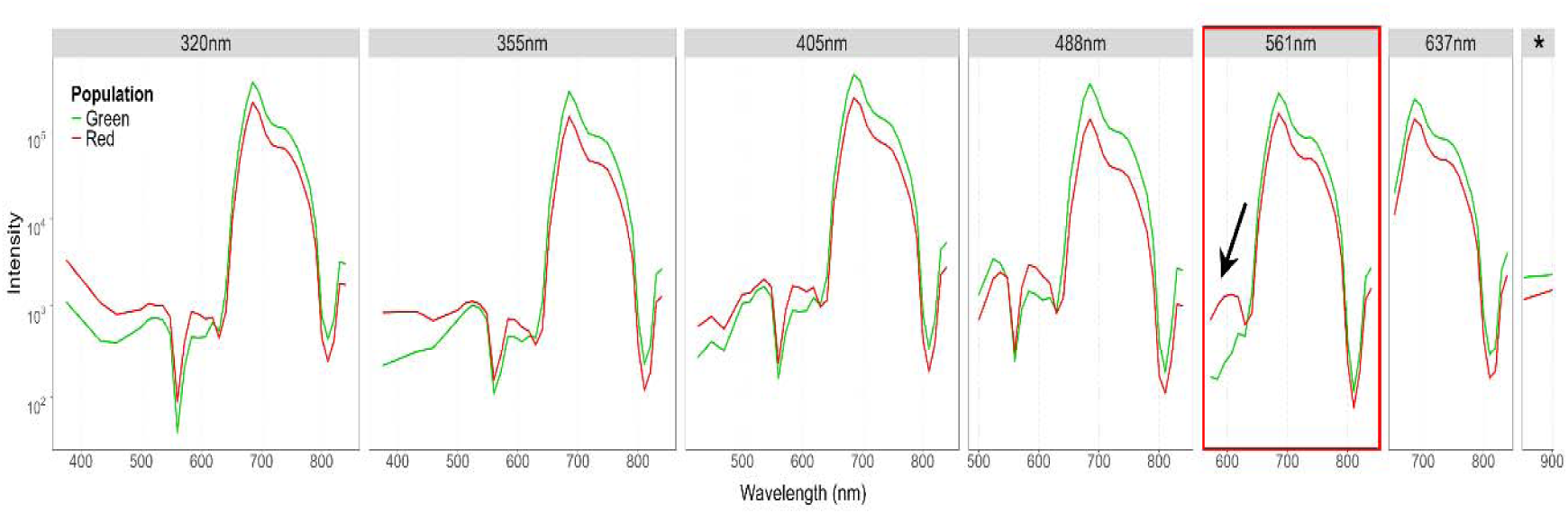
Spectral autofluorescence shifts in normal (green) and stressed (red) *Haematococcus* sp. CCAC 3319 cultures (black arrow indicates accessory pigments region – R2), * - 808 nm excitation laser wavelength.

### Spectral autofluorescence for monitoring pigment biosynthesis and microbial associations

To further investigate the relationship between spectral autofluorescence and physiological state, we analyzed both normal and stressed cultures of *Haematococcus* sp. CCAC 3319, a species known for its dynamic pigment responses to stress (Figure 4). The stressed cultures, characterized by red coloration, exhibited distinct spectral profiles, with an increase in fluorescence intensity across in the ∼566–625 nm range at 561 nm excitation. Although some moderate spectral variation was observed at 355 nm and 405 nm excitation, the most pronounced differences occurred at 561 nm, highlighting this wavelength’s sensitivity to pigment state transitions. In a separate analysis, we looked at the impact of bacterial presence on autofluorescence by comparing xenic and axenic cultures of *Gonium pectorale* (CCAC 0085B and CCAC 0085, respectively). The xenic culture, which includes a natural bacterial community, displayed altered spectral signatures compared to the axenic culture. This suggests that microbial associations or shifts in the host’s metabolic state may influence the autofluorescence properties (Supplementary Figure 5).

Furthermore, to investigate shifts in pigment-related autofluorescence, we monitored *Serratia marcescens* over a 15-hour growth period on solid media. Time-resolved spectral measurements revealed a distinct shift beginning around the 9-hour mark, with significant changes observed under 488 nm and 561 nm excitation. A prominent emission peak developed around ∼570 nm (Figure 5). This shift likely indicates the accumulation of prodigiosin, a tripyrrolic red pigment synthesized by *S*.*marcescens* in response to specific growth conditions^36^ and was accompanied by additional emission in the far-red region. These findings demonstrate that autofluorescence can be a valuable, non-invasive tool for tracking pigment biosynthesis and metabolic changes in algae and bacteria.

**Figure 5.**
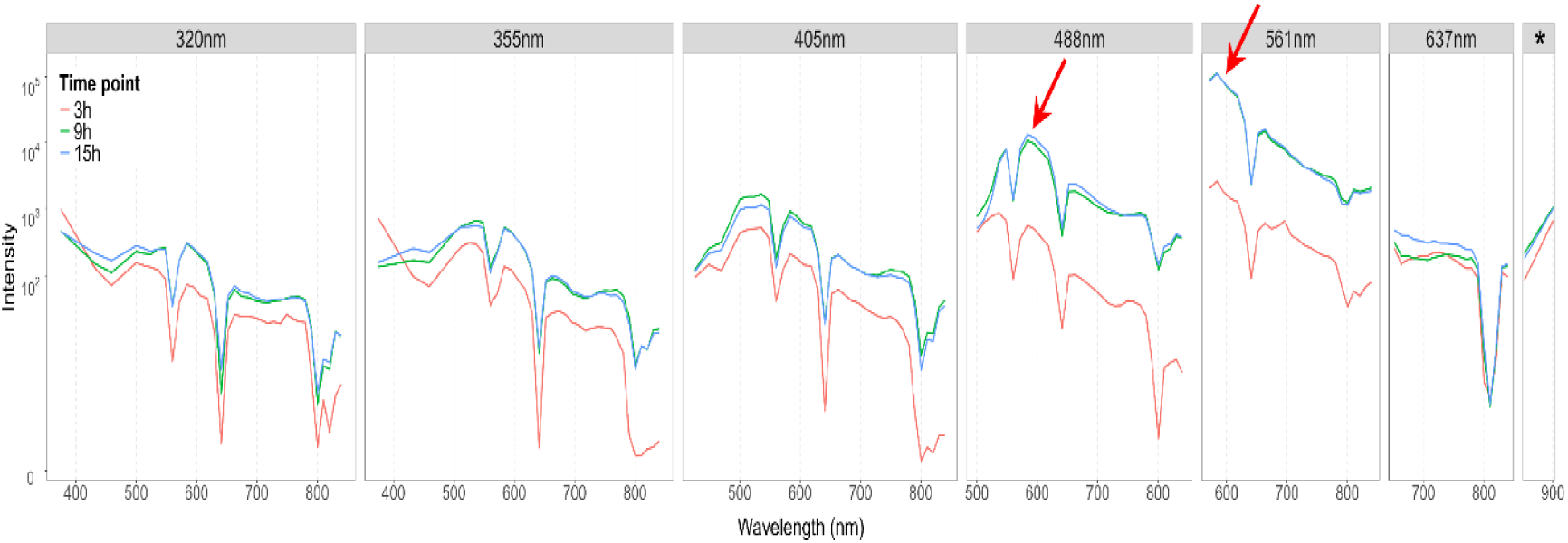
Time-course spectral analysis of red-pigmented *S.marcescens* (grown on solid media over 15 hours); red arrows indicated emission peaks of interest.

## Discussion

Microalgae are classified into different size groups, and their morphology is typically examined using light and fluorescent microscopy. Variations in pigments differences not only help cells to adapt to environmental conditions but can also serve as markers for taxa classification based on their unique pigment profiles^15^. For instance, the presence of fucoxanthin in diatoms, peridinin in dinoflagellates, and phycobiliproteins in cyanobacteria and cryptophytes helps differentiate these groups based on their pigment composition^37^. Pigments play a crucial role in algal physiology, driving photosynthesis, photoprotection, and stress responses^4^. Additionally, other substances that are present during the analysis of microalgae, such as detritus and dissolved organic matter^38^, can also exhibit fluorescence.

Algal pigment analysis has traditionally relied on high-performance liquid chromatography (HPLC) because of its high specificity and sensitivity^17^. HPLC provides detailed pigment profiles that are useful for chemotaxonomy and phytoplankton community analysis by detecting chlorophylls, carotenoids, and their degradation products^17^. Other techniques, such as spectrophotometry, fluorescence microscopy, and remote sensing, serve as alternative options for pigment analysis^39^. However, similar to HPLC, these methods also depend on population average measurements ^20,40^. In contrast, full-spectrum cytometry has emerged as a high-throughput, single-cell alternative. providing information about cell size and scattering while capturing full autofluorescence emission spectra across multiple excitation wavelengths (320-808 nm for ID7000, Sony Biotechnology, Fremont, CA, USA) ^28,29,41,42^.

In this study, we analyzed the autofluorescence properties of nine major microalgal groups to determine whether distinct spectral differences could be detected at the phylum level. We found that chlorophyll-based fluorescence alone is insufficient for taxonomic discrimination^43^. However, certain algae, such as *Cryptomonas* spp., exhibited a unique Chl *a*_682_ emission peak at physiological temperatures^44^, which we also observed in our full-spectrum emission data. Most algal phyla were distinguished by variations in the fluorescence of accessory pigments, particularly in the green and yellow-orange regions. Our findings are consistent with previous studies on PBP- and no-PBP-containing algae, which reported emission wavelength peaks of phycoerythrin at 570 to 600 nm and phycocyanin at 650 nm consistent with the peaks observed in PBP-containing taxa^45^. There were some differences in the fluorescence emission of phycobilin pigments across various algal groups, which were reported to range from 560 to 578 nm for *Synechococcus* spp.^32,33^, and from 578 to 617 nm for Cryptophyta^45–47^. Additionally, our results demonstrated that the choice of excitation wavelength significantly impacts spectral differentiation. As mentioned previously, the diversity of pigments in algae is shaped by evolutionary processes, particularly through primary and secondary endosymbiosis^48^. The results revealed the emergence of distinct algal lineages, each with unique pigment compositions, including some PBP-containing taxa^49^.

The seemingly uniform algal populations exhibit spectral and morphological diversity that varies based on their growth stage and whether they are under normal or stress conditions. An analysis of a single isolate of *Chlamydomonas* sp. revealed five distinct autofluorescent subpopulations that differ in Chl *a* intensity, cell size and shape. This finding indicates that cell morphology significantly influences autofluorescence spectral properties. Alternatively, variations in pigment organization within the cell may also contribute to this spectral variability. Furthermore, imaging cytometry-based analysis confirmed the existence of different subpopulations with shifted chlorophyll autofluorescence. Together, these findings demonstrate that autofluorescence can serve not only as a taxonomic marker but also as an indicator of physiological cellular state and metabolic heterogeneity within cultures. The presence of various algal subpopulations, including both actively growing and dying cells, in monocultures necessitates a combination of scattering and fluorescence analysis, which cannot be achieved through traditional population averaging methods.

Chl *a* concentration is widely used for the indirect determination of algal biomass across various algal groups using methods such as spectrophotometry, fluorometry, HPLC, and, more recently, conventional flow cytometry^50–52^. Our findings, consistent with previous studies, indicate that Chl *a*-based red autofluorescence can persist long after cell death, which makes it inadequate for viability assessments^43^. By using Chl *a* fluorescence as a proxy for cell sorting, we were able to identify different algal populations and their unique spectral signatures. While yellow-orange fluorescence is well-characterized in taxa containing PBPs, its role in algae lacking these pigments remains uncertain. Prior research has shown that exposure to light, nutrient stress, and oxidative damage can trigger the synthesis of secondary carotenoids as a mechanism for photoprotection and cellular defense^53,54^.

Differences in metabolic cell activities are linked to specific fluorescence regions of the spectrum, particularly in the green and yellow-orange ranges. An increase in green autofluorescence (GF) has been observed following cell death or fixation, which may indicate changes in pigment composition or responses to oxidative stress responses^55^. While the molecular basis of green autofluorescence remains poorly understood, studies suggest that it may originate from flavin-containing proteins^56–60^. Additionally, GF may be associated with stress responses, alterations in pigments, and small molecules such as NADH and FAD^55^, which have been linked in cells to metabolic activity of flavins. NADH and FAD are associated with green fluorescence across both prokaryotic and eukaryotic cells, with NADH fluorescence often linked to active respiration and metabolic activity^61,62^. By integrating assessments of the green and yellow-orange fluorescence spectral regions, we can improve our ability to identify metabolic and morphological differences among microalgal populations.

The application of spectral cytometry extends beyond the study of microalgae and into the realm of microbiological analysis. The pigmented bacterium *Serratia marcescens* has exhibited unique spectral signatures and shows pigment-dependent shifts in autofluorescence during its growth. This is further supported by research on prodigiosin (PG), a red tripyrrole pigment with pH-dependent spectral properties^63,64^. As a result, the recorded variations in autofluorescence observed in *Serratia* cultures may indicate changes in PG biosynthesis, providing a potential non-invasive method for monitoring microbial pigment production in biotechnological processes.

This work demonstrates that full-spectrum cytometry can enhance our ability to catalog and organize the inherent autofluorescent heterogeneity of microalgae under various environmental conditions. By utilizing autofluorescence-based spectral cytometry, we can differentiate various taxa based on the pigment composition of microalgae and microbes, particularly in spectral regions associated with PBPs and carotenoids. This method links variations in fluorescence intensity, the shape of chlorophyll peaks, and the spectral regions associated with accessory pigments to cellular diversity and physiological states of microalgae. The combination of autofluorescence-based sorting and IFC has shown that cells with different morphological characteristics such as shape and size, exhibit distinct spectral signatures. Expanding this approach beyond microalgae, research on bacteria, such as *S.marcescens*, has revealed growth-dependent shifts in pigment composition, highlighting the broader applicability of spectral cytometry in microbial ecology. Together these findings position autofluorescence-based analysis as a high-throughput method for taxonomic differentiation, cellular profiling, and stress detection in microorganisms.

Future research should focus to broaden spectral libraries for different microalgal and bacterial taxa in order to improve classification accuracy and uncover new insights into pigment-based spectral variations. Additionally, integrating spectral cytometry with single-cell imaging and cell sorting has the potential to significantly advance biotechnology and algal research.

## Institutional Review Board Statement

Not applicable.

## Informed Consent Statement

Not applicable.

## Data Availability Statement

Detailed data are available from corresponding author by reasonable request.

## Acknowledgments

We are acknowledging excellent help from Vladimir Novokhatsky, Nazarbayev University, Steffen Boehner, Rudolf Bichele, Vendula Sinkorova from Sony Biotechnologies (USA), Grigory Marchenko, Cytek-Amnis (USA), and Aigul Kussanova from Core Facilities, Nazarbayev University.

